# Non-canonical Sodium Channel Isoforms Underlie Chamber Specific Cardiac Excitability

**DOI:** 10.1101/2025.08.27.672688

**Authors:** Colin J. Clark, Christian Anderson, Alex Dou, Jason Dierdorff, Jason D. Galpin, Lionel Gissot, Samantha Thompson, Hannah Choi, Jin-Young Yoon, Daniel T. Infield, Jared McLendon, Peter Bronk, Keane Leeds, Ryan L. Boudreau, Bum-Rak Choi, Barry London, Christopher A. Ahern

## Abstract

Voltage-gated sodium (Na_V_) channels drive cardiac excitability. While Na_V_1.5 is the primary cardiac isoform, the composition and functional contributions of non-Na_V_1.5 isoforms in the heart remain unclear. Here, we developed a chemical-genetic mouse model (Na_V_1.5-GX) in which Na_V_1.5 can be selectively and reversibly inhibited by acyl- and aryl-sulfonamide compounds (GX drugs). Na_V_1.5-GX mice exhibited normal cardiac function at baseline, but acute GX drug administration caused profound conduction defects and arrhythmias. Whole-heart optical mapping revealed dose-dependent chamber-specific sensitivity to Na_V_1.5 inhibition, with the right ventricle (RV) being the most sensitive, followed by the left ventricle (LV), left atrium (LA), and right atrium (RA). Patch-clamp recordings of isolated cardiomyocytes with application of Na_V_ isoform-selective inhibitors showed that Na_V_1.5 contributed 93% of sodium current in the LV, 81% in the RV and 78% in the LA. Non-Na_V_1.5 isoforms were differentially enriched across chambers: Na_V_1.8 in the LV, Na_V_1.1/1.3 in the RV, and Na_V_1.2/1.6/1.7 in the atria. These results reveal a surprising chamber-specific isoform landscape of cardiac sodium currents which may underlie the right ventricular predominant phenotype of Brugada syndrome and highlight non-Na_V_1.5 isoforms as potential mediators of chamber-specific cardiac pathologies and as pharmacological targets.

## Introduction

Cardiac arrhythmias are common clinical conditions resulting from disruption of normal electrical activity in the heart. They may be either inherited or acquired and occur in either the atria or the ventricles, with the most common arrhythmia being atrial fibrillation. Ventricular arrhythmias are less common but of particular interest due to their high mortality and association with other disease states. Treatment often involves the use of antiarrhythmic medications which exert their effects through modulation of the cardiac action potential. Vaughan Williams Class I medications are among the most frequently prescribed drugs and work via inhibition of the inward sodium current in the heart.^1,2^

Voltage-gated sodium (Na_V_) channels drive the rapid upstroke of the action potential in cardiomyocytes and enable synchronous electrical conduction through the heart. The predominant cardiac isoform, Na_V_1.5 (encoded by *SCN5A*), generates the bulk of inward sodium current (I_Na_) and is a major target for class I antiarrhythmic drugs, such as flecainide. Mutations in *SCN5A* are implicated in a broad spectrum of cardiac disorders, including Brugada syndrome, long QT syndrome, conduction disease and atrial fibrillation,^3^ reflecting the clinical importance of Na_V_1.5 to cardiac function. However, chamber-specific electrical phenotypes, such as the right ventricular predominance of the Brugada pattern on electrocardiography,^4^ and increased atrial sensitivity to flecainide,^5^ suggest that Na_V_1.5 alone may not account for cardiac action potential generation in each heart chamber.

Although Na_V_1.5 is recognized as the primary source of cardiac I_Na_, it has become generally accepted that non-Na_V_1.5 isoforms, typically considered neuronal, are present in the heart. These may include Na_V_1.1, Na_V_1.2, Na_V_1.3, Na_V_1.4, Na_V_1.6, and Na_V_1.8, which have been detected at the transcript and protein levels in both humans and animal models.^6–11^ The presence of these channels also appears to be different between the atria and the ventricles.^10^ However, the functional relevance of these isoforms in the heart remains poorly understood. Existing pharmacological tools and genetic models lack the specificity required to selectively and completely suppress Na_V_1.5, limiting the ability to parse isoform-specific contributions.

To address this, we developed a chemical-genetic mouse model in which the Na_V_1.5 isoform is rendered sensitive to selective pharmacologic blockade by acyl- and aryl-sulfonamide compounds (GX drugs),^12,13^ originally developed to inhibit Na_V_1.7 for pain treatment. This approach enables functional isolation of Na_V_1.5 from other Na_V_ isoforms in native cardiomyocytes. We combined this model with in vivo electrophysiology, ex vivo optical mapping, and whole-cell patch clamp to systematically map the composition and chamber-specific contribution of cardiac Na_V_ isoforms.

Our findings reveal that each cardiac chamber exhibits a distinct pattern of Na_V_ isoform usage, with non-Na_V_1.5 isoforms playing significant, regionally specific roles in supporting cardiac excitability. These results provide a molecular framework for understanding chamber-specific arrhythmogenesis as seen in Brugada syndrome. It also provides valuable insight into potential therapeutics and possible off-target effects for the growing number of small molecule drugs designed to specifically target non-cardiac isoforms, such as Na_V_1.8,^14^ previously not known to play a clear role in cardiac excitation and conduction.

## Results

### Engineered selective inhibition of cardiac Na_V_1.5 by GX drugs

To enable selective pharmacologic control of Na_V_1.5 current, we genetically engineered a Na_V_1.5 channel construct with sensitivity to GX drugs, known as the Na_V_1.5-GX channel. This was achieved by introducing the GX binding site in the Domain IV S2-3 region of Na_V_1.7 (Q1556-V1620) into the Na_V_1.5 channel (P1556-F1620) background (**Fig. 1A**) via CRISPR-Cas9 editing of the native *SCN5A* locus (exons 27 and 28) in C57BL/6J mice (**Fig. S1**). The resulting chimeric allele (Na_V_1.5^GX^) exhibited simple Mendelian inheritance and normal cardiac development, including normal electrocardiography (ECG) and echocardiography measurements over the first 12 months of life (**Fig. S2**). ECG and echocardiography showed time dependent decreases in LVEF, with increases in HR and decreases in QTc interval at 12 months likely attributable to sedation during ECG acquisition. Importantly, there were no differences when comparing between groups at a single time point.

**Figure 1.**
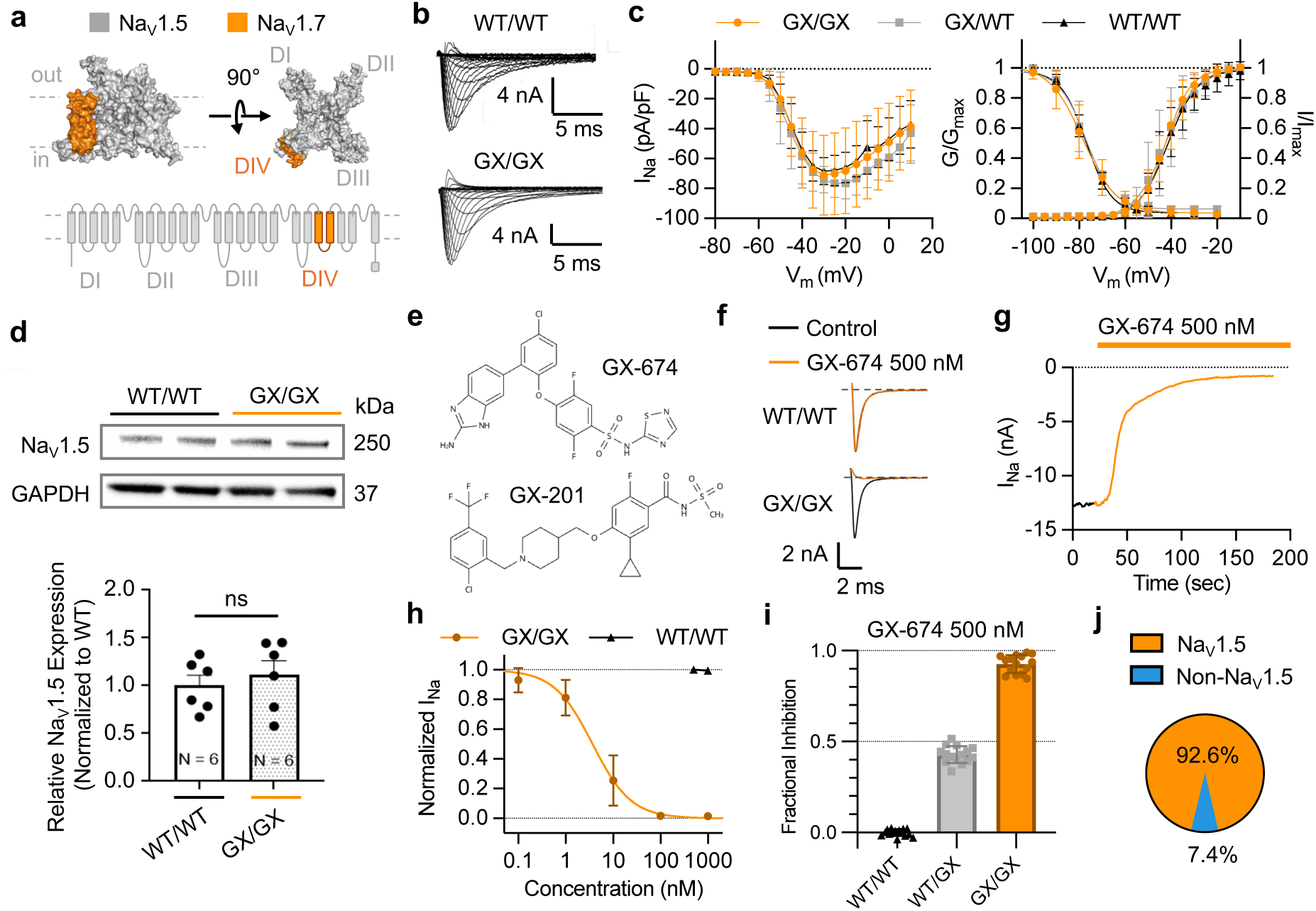
Engineered GX drug sensitivity enables selective inhibition of NaV1.5 current in mouse cardiomyocytes. (A) Representative structure (top) and topology (bottom) of the NaV1.5-GX chimeric channel (rat NaV1.5, PDB ID: 7FBS). The GX binding site in S2-3 of the NaV1.7 DIV voltage sensor was introduced into the NaV1.5 background via CRISPR-Cas9 editing of the *SCN5A* locus in C57BL/6J mice. (B) Representative whole-cell sodium current tracings in left ventricular cardiomyocytes from WT (Nav1.5^+/+^, top) or Nav1.5-GX (Nav1.5^GX/GX^, bottom) mice. (C) Current-voltage (left), conductance-voltage (right, left axis), and steady-state inactivation relationships (right, right axis) from left ventricular cardiomyocytes show no differences in WT (Nav1.5^+/+^, black), heterozygous (Nav1.5^+/GX^, gray), or homozygous NaV1.5-GX (Nav1.5^GX/GX^, orange) mice. (D) Western blots (top) with relative protein quantification normalized to GAPDH (bottom) show no difference in Nav1.5 protein expression in whole cell lysates from WT or homozygous NaV1.5-GX ventricular cardiomyocytes. (E) Structure of the inhibitory GX drugs GX-674 (top) and GX-201 (bottom). (F) Representative whole-cell sodium current traces from WT (top) and NaV1.5-GX (bottom) left ventricular cardiomyocytes before (black) and after (orange) addition of 500 nM GX-674. (G) Representative diary plot of peak currents during addition of 500 nM GX-674 (orange bar). Cells were held at -80 mV, hyperpolarized to -120 mV for 100 ms, and depolarized to -20 mV for 300 ms to elicit peak currents measured at 1 Hz in 50 mM Na^+^ external solution. (H) Dose-response curve of GX-674 on WT (black, triangles) and NaV1.5-GX (orange, circles) left ventricular cardiomyocytes. (I) Maximal GX-674 inhibition in WT (Nav1.5^+/+^, black), heterozygous (Nav1.5^+/GX^, gray), or homozygous NaV1.5-GX (Nav1.5^GX/GX^, orange) left ventricular cardiomyocytes. (J) Proportion of the sodium current attributable to Nav1.5 (orange) or non-Nav1.5 isoforms (blue). Mean ± SD. Statistical significance determined by Student’s t-test (α=0.05).

To determine if introduction of the GX-binding site disrupts Na_V_1.5 channel function, we performed whole-cell patch clamp electrophysiology of sodium currents in isolated left ventricular cardiomyocytes from Na_V_1.5^+/+^ (WT), Na_V_1.5^+/GX^, and Na_V_1.5^GX/GX^ mice. Cardiomyocytes expressing Na_V_1.5-GX showed no difference in peak sodium current density, voltage-dependence of activation, or steady-state inactivation relative to cardiomyocytes expressing WT Na_V_1.5 in the absence of GX drug (**Fig. 1B-C**). RNA sequencing (RNA-Seq) of bulk murine left and right ventricles showed no significant differences in voltage gated sodium channel subunit between Na_V_1.5^+/+^ and Na_V_1.5^GX/GX^ animals in the absence of GX drug (**Table S1**). Likewise, western blots of ventricular cardiomyocyte whole-cell lysates showed no difference in baseline Na_V_1.5 protein levels between Na_V_1.5^+/+^ and Na_V_1.5^GX/GX^ mice (**Fig. 1D**). Hence, the Na_V_1.5-GX channel functionally recapitulates WT Na_V_1.5 at baseline.

Next, we assessed the ability of GX drugs (**Fig. 1E**) to selectively bind and inhibit the Na_V_1.5-GX channel by measuring cardiac sodium current responses to GX-674, a compound developed to selectively inhibit Na_V_1.7. GX-674 robustly inhibited the sodium current in left ventricular cardiomyocytes of Na_V_1.5-GX mice at nanomolar concentrations (IC50 = 3.05 ± 0.08 nM), whereas that of WT mice showed no response to the drug up to 1 μM (**Fig. 1F-H**). Importantly, a saturating dose of GX-674 (500 nM) inhibited less than half (42.8 ± 4.6%) of the sodium current in Na_V_1.5^+/GX^ mice and most but not all (92.6 ± 4.9%) of the current in Na_V_1.5^GX/GX^ mice (**Fig. 1I**). The observed residual GX-resistant current constitutes 7.4 ± 4.9% of the baseline current in left ventricular myocytes, suggesting the presence of functional non-Na_V_1.5 isoforms in cardiomyocytes (**Fig. 1J)**.

### Acute inhibition of Na**_V_**1.5 induces arrhythmias in vivo

To test the effects of acute Na_V_1.5 suppression in vivo, we used surface ECG to monitor cardiac conduction before and after intraperitoneal (IP) or oral administration of GX drugs. IP administration of GX-201 caused dose dependent decreases in heart rate (HR) as well as PR and QRS prolongation measured one hour after drug administration in Na_V_1.5-GX but not WT mice (**Fig 2A-D**). At 5 mg/kg, the average HR was 433.3 ± 169.8 BPM (574.7 ± 49.3 BPM in WT, p<0.05), PR interval was 66.33 ± 14.64 ms (40.01 ± 3.16 ms in WT, p<0.0001), and QRS duration was 45.34 ± 12.23 ms (15.17 ± 0.98 ms in WT, p<0.0001). IP administration at doses greater than 2.5 mg/kg resulted in atrioventricular block in 3 of 6 mice (**Fig. 2M**), and ventricular tachycardia in 5 of 6 mice (**Fig. 2N**).

**Figure 2.**
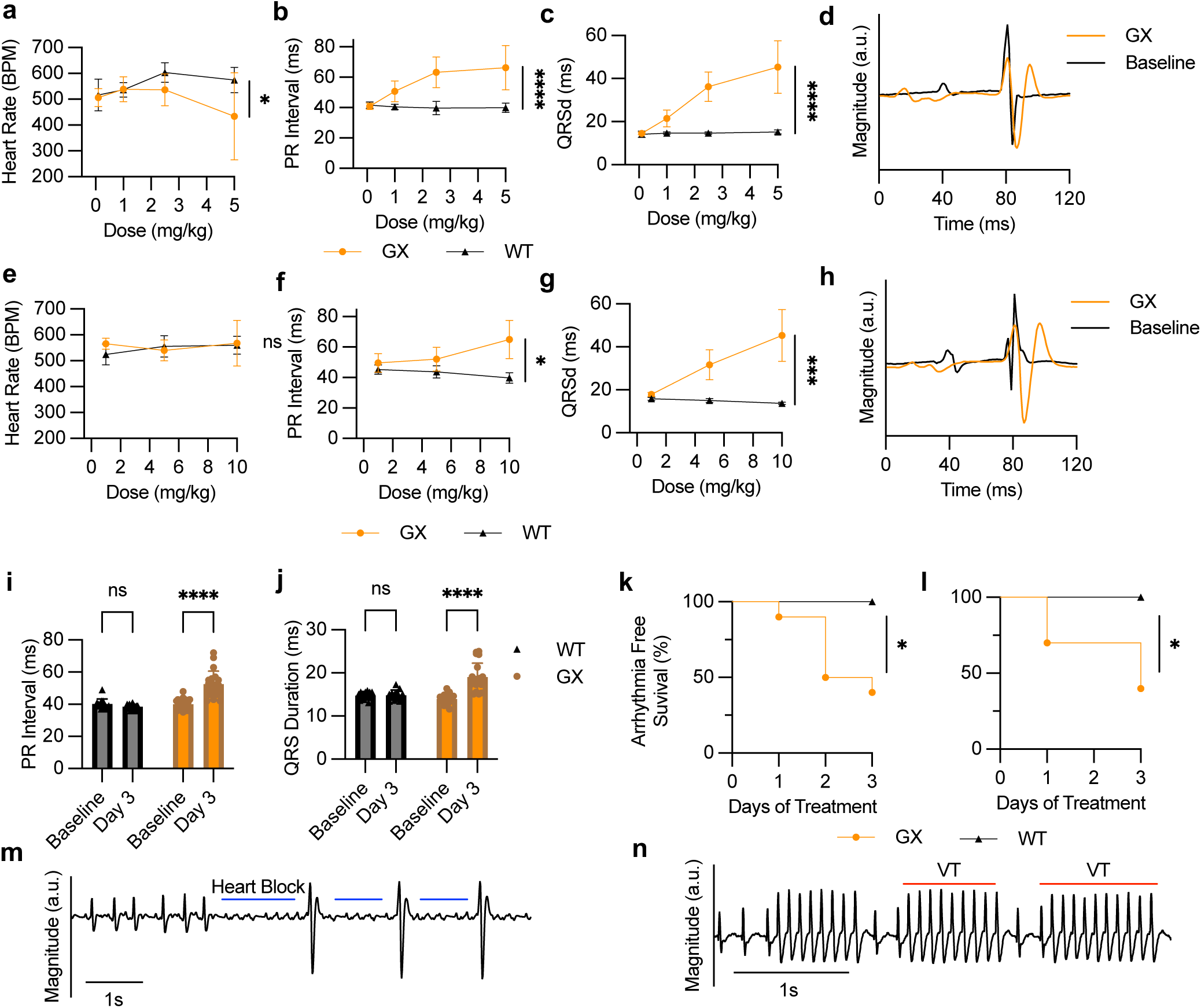
Selective NaV1.5 inhibition induces conduction defects and cardiac arrhythmias in vivo. (A-H) Cardiac electrophysiology meters of WT (black, triangles) and NaV1.5-GX mice (orange, circles) after intraperitoneal (A-D) or oral (E-H) administration of GX-201. Heart (A, E), PR interval (B, F), QRS duration (C, G), and representative traces in lead modified V (D, H) were measured by ECG 1 hour after IP inistration and after 3 days of oral administration of GX-201. (I-J) Comparison of PR interval (I) and QRS duration (J) of WT (black, triangles) Nav1.5-GX mice (orange, circles) before and after 3 days of 2.5 mg/kg/day oral GX-201. (K-L) Kaplan-Meier curves of WT (black, triangles) aV1.5-GX (orange, circles) mice without bradyarrhythmia (K) or tachyarrhythmia (L) on ECG telemetry over 3 days of 2.5 mg/kg/day oral GX-Bradyarrhythmia status was defined as >2.5% of R-R intervals longer than the longest 1% of R-R intervals at baseline, and tachyarrhythmia us was defined as >0.5% of total R-R intervals shorter than the 0.1% shortest R-R intervals at baseline. (M-N) Example ECG traces of heart k (blue lines, M) and non-sustained ventricular tachycardia (VT, red lines, N) and after 5 mg/kg IP injection. Mean ± SD. Statistical significance rmined by two-way ANOVA (I-J) or log-rank test (K-L) at α=0.05 (*, p<0.05; **, p<0.01; ***, p<0.001, ****, p<0.0001). N>3 for all plots.

Similarly, a 3-day treatment with oral GX-201 resulted in dose dependent increases in PR interval and QRS duration in Na_V_1.5-GX but not WT mice (**Fig. 2E-H**). At an oral dose of 10 mg/kg, the average PR interval was 65.00 ± 12.49 ms (39.75 +/- 3.40 ms in WT, p<0.05) and QRS duration was 45.33 ± 11.99 ms (13.75 ± 0.50 ms in WT, p<0.001). Notably, a lower dose of 2.5 mg/kg was sufficient to increase PR interval to 52.50 ± 8.17 ms (38.46 ± 1.71 ms in WT, p<0.0001) and QRS duration to 19.05 ± 3.25 ms (14.85 +/- 1.18 ms in WT, p<0.0001) after 3 days in a larger cohort (**Fig. 2I-J**). Unlike IP administration, heart rate showed no change in response to treatment with oral GX-201. Hence, administration of parenteral or oral GX-201 disrupts normal cardiac electrical activity in a dose- and time-dependent fashion in Na_V_1.5-GX mice but not WT mice, consistent with selective Na_V_1.5 inhibition.

To determine if selective Na_V_1.5 inhibition can induce arrhythmias, we performed 24-hour ECG radiotelemetry on ambulatory mice during 3 days of treatment with a 2.5 mg/kg/day GX-201 diet, followed by a 3-day washout period with a control diet. After 3 days of GX-201 administration, average serum drug levels were 928 ± 591 nM (**Fig. S3**). To quantify arrhythmia burden, bradycardic and tachycardic beats were identified from R-R interval histograms (**Fig. S4**) as R-R intervals in the longest 1.0% or shortest 0.1% of beats at baseline, respectively, and arrhythmias were confirmed by direct review of the tracings. By day three of exposure, over 50% of the Na_V_1.5-GX mice had developed recurrent bradycardia (**Fig. 2K**) or tachycardia (**Fig. 2L**), whereas none of the WT mice showed recurrent arrhythmias. In Na_V_1.5-GX mice, arrhythmias resolved after 24 hours of washout. Of note, histograms of R-R intervals showed development of a population of short R-R intervals corresponding to ventricular tachyarrhythmias (**Fig. S4,** red arrows) and loss of the typical bimodal distribution of R-R intervals corresponding to circadian activity, which returned to baseline after drug washout. Hence, acute suppression of Na_V_1.5 current via oral GX-201 induces reversible arrhythmias in Na_V_1.5-GX mice and therefore models the effects of selective Na_V_1.5 inhibition in vivo.

To evaluate whether GX drug-mediated Na_V_1.5 inhibition causes acute remodeling of gene expression for other primary regulators of cardiac conduction, we performed mRNA-seq to assess cardiac transcriptomic changes in Na_V_1.5-GX and WT mice 4 hours after treatment with the GX drug MRL5^12,15^ (**Supplemental Data Table 1**). This analysis revealed that acute Na_V_1.5 inhibition in Na_V_1.5-GX mice does not result in significant myocardial gene expression changes compared to vehicle-treated littermate controls (based on adjusted q-value <0.05). To evaluate for possible subtle effects, we performed a targeted assessment of cardiac conduction-related genes (based on gene ontology annotations), which indicated only subtle dysregulation in four genes (*Agt*, *Kcne4*, *Atp2b3*, *Nkx2-5*; unadjusted p<0.05). Indeed, most conduction-related genes did not show significant (nor trending) expression changes within the short timeframe tested, supporting the specificity of this model for early assessments of cardiac electrical changes after acute Na_V_1.5 inhibition (**Fig. S5)**.

### Na_V_1.5 inhibition causes chamber-specific changes in cardiac excitability

To assess the effect of selective Na_V_1.5 inhibition on regional cardiac conduction, we performed optical mapping of ex vivo perfused WT and Na_V_1.5-GX hearts using a voltage-sensitive fluorescent dye to visualize whole-heart action potential propagation (**Fig. S6, Supplemental Video 1)**.

At baseline, hearts from WT and Na_V_1.5-GX mice showed no difference in electrical conduction, including longitudinal and transverse conduction velocities or action potential durations in left and right ventricle (**Fig. 3A-B**). Application of increasing doses of GX-674 decreased ventricular excitability in response to pacing and produced regional conduction blocks in Na_V_1.5-GX but not WT hearts (**Fig. 3C, Supplemental Video 2**). Conduction deficits in Na_V_1.5-GX hearts corresponded to a decrease in longitudinal and transverse conduction velocity and increases in action potential rise time in left and right ventricle relative to WT hearts (**Fig. 3D**).

**Figure 3.**
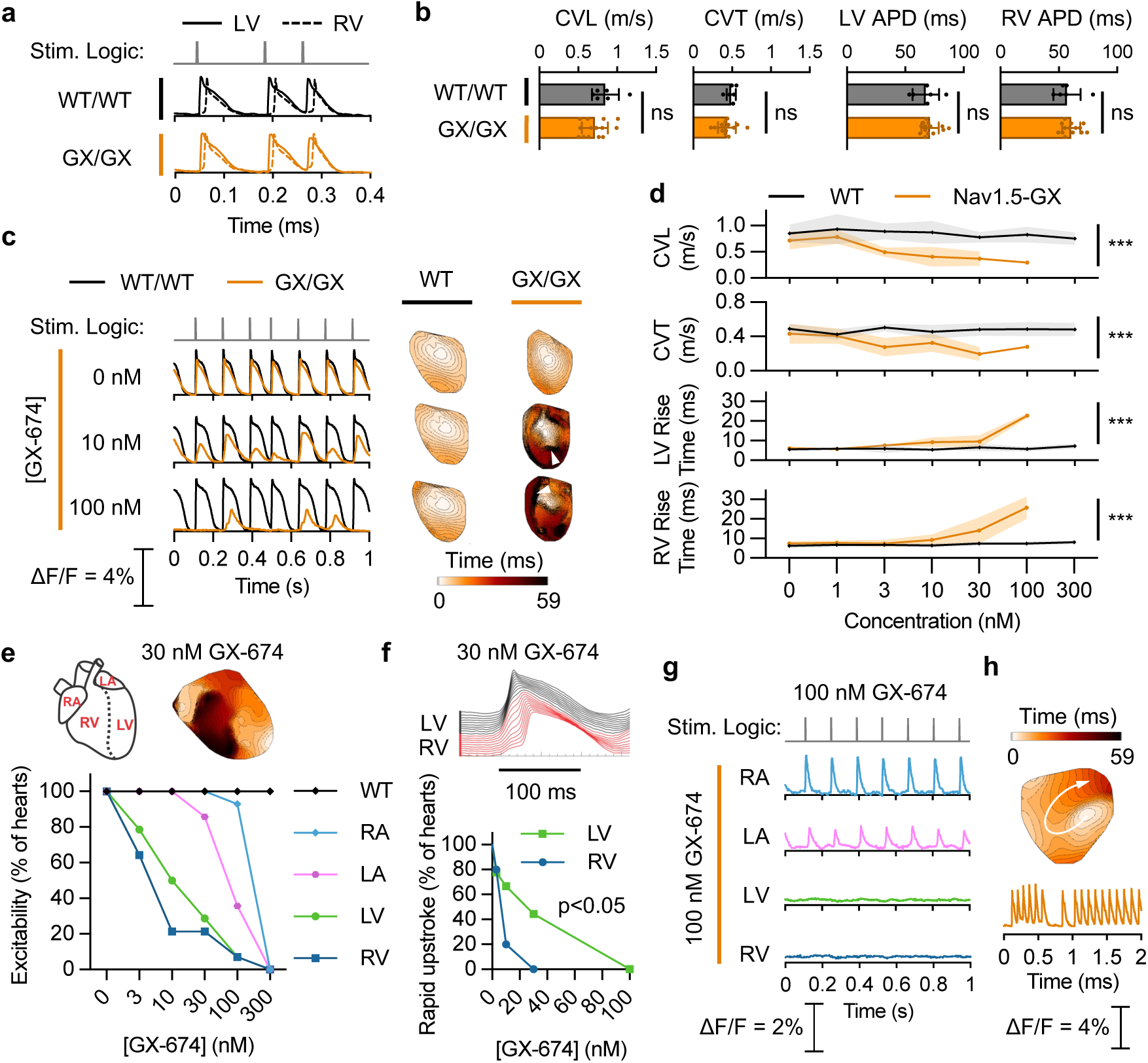
NaV1.5 inhibition by GX drugs disrupts cardiac excitability in a chamber-specific manner. (A) Baseline left (solid line) and right (dashed line) ventricular action potentials are similar in NaV1.5-GX (orange) and WT (black) mice. (B) The GX allele does not change the baseline cardiac conduction velocity (longitudinal, CVL; transverse, CVT) or action potential duration (APD). Mean ± SD; n=14 GX, 6 WT hearts; p>0.05 (ns), independent t-test. (C) GX-674 reduces cardiac excitability in NaV1.5-GX but not WT hearts. Sample LV AP traces at 0, 10, and 100 nM GX-674 (left) and activation maps (right) showing iso-temporal lines from the stimulus site (lightest color). Areas of conduction block are indicated (white triangles). (D) GX-674 slows ventricular conduction velocity and increases rise time in a dose dependent manner. Mean ± SD; ***, p<0.001; two-way ANOVA linear model with Bonferroni correction. (E) The response of NaV1.5-GX hearts to GX-674 is heterogeneous between cardiac chambers. Schematic (top left) and representative activation map (top right, scale as in C) of a NaV1.5-GX heart paced at the right atrium in the presence of 30 nM GX-674. Dose-response curves (bottom) showing highest sensitivity in the RV, followed by LV; atrial tissues were significantly more resistant (RV vs. RA: p=1.9e-5; LV vs. LA: p=0.02; RA vs. LA: p=0.02; Kolmogorov-Smirnov test). (F) Conduction block was characterized by biphasic and delayed action potential upstrokes (top; space-time plots from E; RV, red; LV, black) which occur at lower GX-674 concentrations in the RV than the LV (bottom; RV, blue; LV, green; p=0.03, log-rank test). (G) Sample AP traces showing chamber-specific changes in excitability of NaV1.5-GX mouse hearts in the presence of 100 nM GX-674. (H) NaV1.5-GX hearts showed spontaneous ventricular tachycardia (n=4/16 hearts) in the presence of GX-674. Sample activation map (top) showing reentry and action potential traces (bottom).

Strikingly, each heart chamber showed differential sensitivity to Na_V_1.5 inhibition. The dose required to block excitability in >50% of hearts was 10 nM for the right ventricle (RV), 30 nM for the left ventricle (LV), 100 nM for the left atrium (LA), and 300 nM for the right atrium (RA); (**Fig. 3E, Supplemental Video 3**). Loss of excitability in the ventricles was preceded by development of a biphasic action potential upstroke, characterized by a weak early depolarization followed by a larger delayed upstroke. This change in action potential morphology appeared at lower concentrations in the RV than the LV (**Fig. 3F**). At 100 nM GX-674, there was complete loss of electrical excitability in the ventricles but not the atria (**Fig. 3G**).

Application of GX-674 to Na_V_1.5-GX hearts produced episodes of spontaneous ventricular tachycardia arising from ventricular re-entrant loops (**Fig. 3H**), consistent with the tachyarrhythmias observed previously via ECG telemetry. Optical mapping showed areas of conduction slowing and block promoting re-entrant arrhythmias. These arrhythmias occurred at both low (10 nM) and intermediate (30 nM) doses of GX-674 in a dose dependent manner (**Supplemental Video 4**). Together, these data reveal that inhibition of Na_V_1.5 produces chamber-specific disruption of cardiac conduction, suggesting that different regions of the heart exhibit differential reliance on Na_V_1.5 current for their excitability.

### Selective Na_V_ inhibition reveals chamber-specific Na_V_ isoform expression

To define the molecular basis of the observed chamber-specific responses to Na_V_1.5 inhibition, we used isoform-selective Na_V_ toxins and small molecule inhibitors to isolate the contribution of each Na_V_ isoform to the cardiac sodium current in each heart chamber.

Four inhibitors were used at fixed concentrations based on their isoform selectivity, defined as <10% off-target inhibition when applied at a single concentration (**Fig. 4A**): GX-674 at 500 nM (inhibits Na_V_1.2, Na_V_1.6, Na_V_1.7, and Na_V_1.5-GX), μ-conotoxin GIIIB^16^ at 200 nM (inhibits Na_V_1.4), VSTx-3^17^ at 500 nM (inhibits Na_V_1.3, Na_V_1.7, and Na_V_1.8), and TTX at 50 nM (inhibits all isoforms except Na_V_1.5, Na_V_1.8, and Na_V_1.9). Isoform selectivity of GX-674, GIIIB, and VSTx-3 were validated via on-target and off-target inhibition dose-response curves on HEK-293T cell lines expressing Na_V_1.5-GX, Na_V_1.3, Na_V_1.4, Na_V_1.6, and Na_V_1.8 constructs (**Fig. S7**).

**Figure 4.**
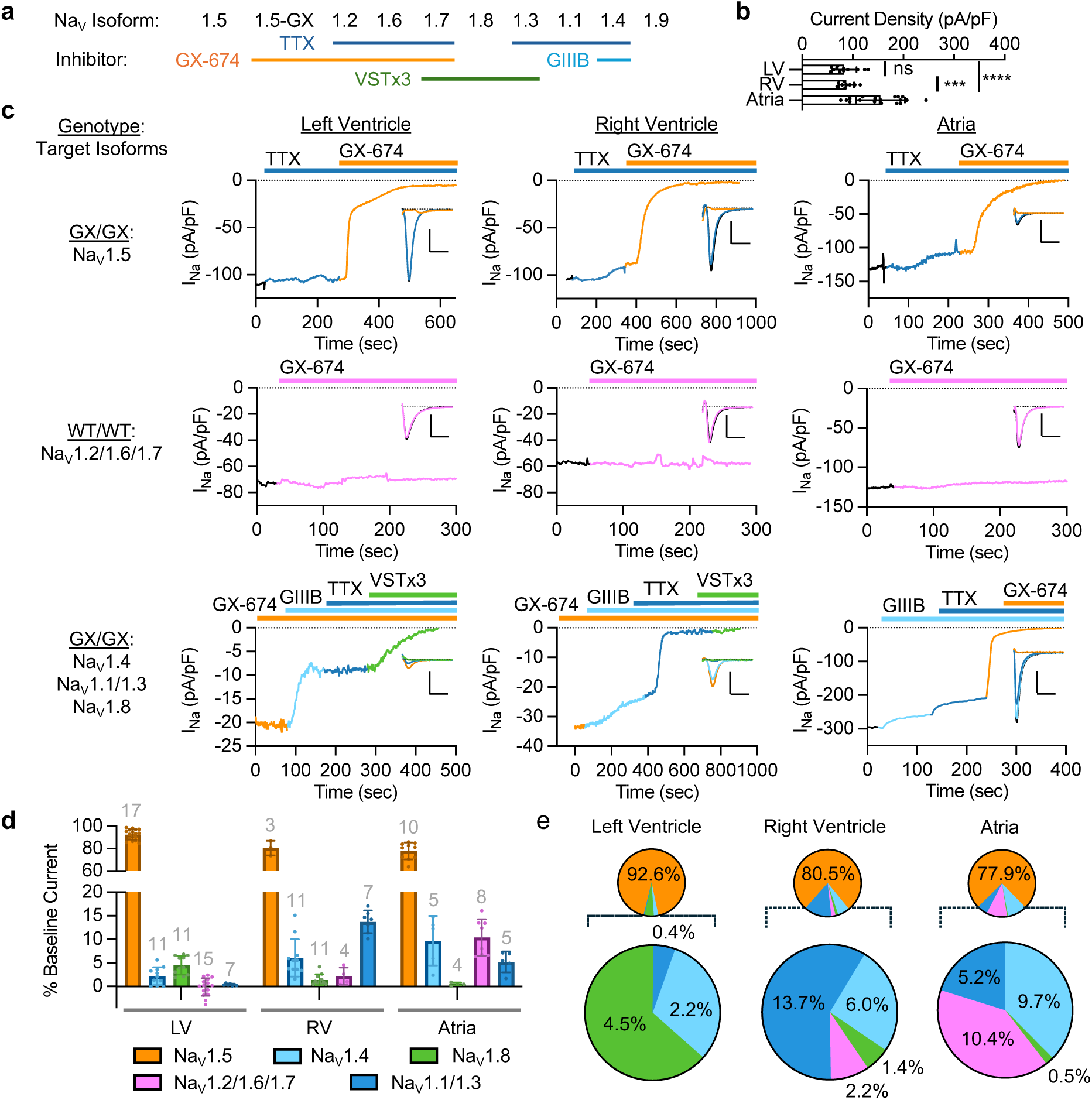
Selective NaV inhibition reveals chamber-specific functional repertoire of NaV isoforms. (A) NaV isoform selectivity of inhibitors GX-674 at 500 nM (orange; NaV1.2/1.6/1.7 and NaV1.5-GX), VSTx3 at 500 nM (green; NaV1.3/1.7/1.8), TTX at 50 nM (dark blue; NaV1.1/1.2/1.3/1.4/1.6/1.7), and μ-Conotoxin GIIIB at 200 nM (light blue; NaV1.4). (B) Average current density is similar in left and right ventricles but higher in the atria. Data from pooled WT and NaV1.5-GX cardiomyocytes (LV, n=13; RV, n=9; atria, n=18). Mean ± SD; ***, p<0.001; ****, p<0.0001; one-way ANOVA with post-hoc Tukey’s test. (C) Representative diary plots of peak current density and current traces (insets) of sodium currents during cumulative serial application of NaV inhibitors in isolated mouse cardiomyocytes from left ventricle (left), right ventricle (middle), and atria (right). Cells were held at a potential of -80 mV, hyperpolarized to -120 mV for 100 ms, and depolarized to -20 mV for 300 ms to elicit peak currents at 1 Hz in 50 mM Na^+^ external solution. Scale bars in insets represent 5 nA (vertical) and 3 ms (horizontal). Bars above each plot indicate the application of NaV inhibitor at the concentrations described above. Experiments targeting NaV1.4, NaV1.1/1.3, and NaV1.8 in ventricle (middle left, bottom left) were performed in 140 mM Na^+^ external solution in the presence of 500 nM GX-674 at baseline to amplify current from the target isoforms. (D) Estimated percent of the baseline sodium current attributable to each NaV isoform in LV, RV, and atria. Percentages of NaV1.4, NaV1.1/1.3, and NaV1.8 were scaled according to the average non-NaV1.5 and non-NaV1.2/1.6/1.7 current. Mean ± SD. Sample sizes are indicated above each bar. N ≥ 6 mice per heart chamber. (E) Summary of sodium current composition by NaV isoform in LV, RV, and atria. Legend as in panel D.

First, we determined total inward sodium current in myocytes isolated from the LV, RV, and LA. There was no significant difference in current density between LV and RV myocytes (83.3 ± 24.1 pA/pF, n=13 vs. 87.6±14.7 pA/pF, n=9, p=0.96) while current density from atrial myocytes was significantly larger (154.6±48.5 pA/pF, n=18, p<0.0001 compared to LV and p<.001 compared to RV; **Fig. 4B**). We then determined the fraction of the baseline sodium current conducted by Na_V_1.5 by applying GX-674 to Nav1.5^GX/GX^ cardiomyocytes. 50 nM TTX was added prior to GX-674 to mask the effects of GX-674 on TTX-sensitive (TTX-S) isoforms (**Fig. 4C, top row**). LV myocytes showed no measurable TTX-S current and were treated with GX-674 alone in most experiments. The Na_V_1.5 component was found to be largest in the LV (92.3 ± 4.6%), followed by the RV (80.5 ± 6.5%) and atria (77.9 ± 7.5%) (**Fig. 4D-E**). To determine the collective contribution from Na_V_1.2/1.6/1.7, we next measured the response to GX-674 in WT cardiomyocytes (**Fig. 4C, middle row**). This component was undetectable in LV, small in RV (2.2 ± 1.9%), and largest in atria (10.4 ± 3.9%) (**Fig. 4D-E**).

Finally, we serially applied GIIIB, TTX, and VSTx-3 to parse the contributions of Na_V_1.4, Na_V_1.1/1.3, and Na_V_1.8, respectively (**Fig. 4C, bottom row**). For ventricular myocytes, the GX-resistant current was isolated and amplified by applying GX-674 in a high-sodium bath (140 mM). In atrial myocytes, GX-resistant current amplification was not necessary, so GX-674 was added after GIIIB and TTX. The overall proportion of each isoform was then scaled based on the proportion of non-Na_V_1.5 and non-Na_V_1.2/1.6/1.7 current for myocytes from each chamber. The Na_V_1.4 fraction was most prominent in the atrial myocytes (9.7 ± 5.3%) but also consistently present in ventricular myocytes (LV: 2.2 ± 1.9%, RV: 6.0 ± 4.0%). Notably, Na_V_1.4 accounted for a similar percentage of the non-Na_V_1.5 current across myocytes from each chamber (LV: 31.1%, RV: 25.9%, atria: 37.5%). Na_V_1.8 constituted the majority of the non-Na_V_1.5 current in LV myocytes (4.5 ± 2.0%) but was minimal in RV myocytes (1.4 ± 1.2%) and atrial myocytes (0.5 ± 0.4%). Finally, the Na_V_1.1/1.3 current was significant in RV myocytes (13.7 ± 2.4%), smaller in atrial myocytes (5.2 ± 2.2%), but nearly absent in LV myocytes (0.4 ± 0.2%) (**Fig. 4D-E**).

Together, these data indicate that each heart chamber exhibits a distinct profile of non-Na_V_1.5 isoforms. The LV is characterized by higher expression of Na_V_1.8, the RV by expression of Na_V_1.1/1.3, and the atria by expression of Na_V_1.2/1.6/1.7 and Na_V_1.4 (**Fig. 4E**). Hence, chamber-specific differences in responses to Na_V_1.5 inhibition may be attributed to chamber-specific patterns of expression of both Na_V_1.5 and secondary Na_V_ isoforms.

## Discussion

### The Na_V_1.5-GX mouse is a physiologic model of selective Na_V_1.5 inhibition

The Na_V_1.5-GX mouse represents a novel platform for studying cardiac electrophysiology by enabling targeted suppression of cardiac Na_V_1.5 sodium current. At baseline, the chimeric Na_V_1.5-GX channel retains biophysical and transcriptional properties identical to WT Na_V_1.5, with normal cardiac structure and function in the absence of GX drug. Application of GX drugs potently and selectively inhibits Na_V_1.5-GX current both in vivo and ex vivo, producing cardiac arrhythmias and conduction deficits. Notably, these effects are acutely reversible and do not induce non-specific transcriptional disruptions in stress or inflammatory pathways. Together, these features establish the Na_V_1.5-GX model as a precise tool for studying the function of Na_V_1.5 and non-canonical sodium channel isoforms in the heart.

### Functional Evidence for Non-canonical Na_V_ Isoforms in the Heart

While substantial evidence suggests that neuronal Na_V_ isoforms are present in the heart, their functional role in cardiac excitability has remained elusive. By first selectively inhibiting Na_V_1.5, we provide direct functional evidence that multiple families of non-canonical neuronal Na_V_ isoforms contribute to cardiac sodium currents. Strikingly, each heart chamber expresses a distinct milieu of sodium channel isoforms in addition to Na_V_1.5: Na_V_1.8 is enriched in the LV, Na_V_1.1/1.3 in the RV, and Na_V_1.2/1.6/1.7 and Na_V_1.4 in the atria. Interestingly, all chambers expressed an appreciable amount of Na_V_1.4, the predominant isoform in skeletal muscle. This chamber-specific distribution suggests that non-Na_V_1.5 sodium channels may contribute to regional differences in cardiac conduction and arrhythmogenesis.

### Chamber-Specific Excitability

Optical mapping demonstrates that acute inhibition of Na_V_1.5 suppresses the excitability of each chamber differently: the atria are the most resistant, followed by the left ventricle, and then the right ventricle. This graded response reflects the relative expression of Na_V_1.5 and the repertoire of non-canonical Na_V_ isoforms in each chamber: Atrial myocytes exhibit the highest sodium current density as well as the highest proportion of non-Na_V_1.5 isoforms, potentially providing an increased conduction reserve. In contrast, ventricular myocytes possess a lower sodium current density and increased reliance on Na_V_1.5 for conduction. Notably, early conduction failure in the RV was associated with a slowed and biphasic action potential, which may represent a delay between activation of Na_V_1.5 and other isoforms. The unique sensitivity of the RV to Na_V_1.5 blockade may reflect its smaller total Na_V_1.5 current density (enhancing pharmacologic inhibition) and secondary dependence on Na_V_1.1/1.3, which may be less able to drive action potential generation than Na_V_1.8 in the LV.

This phenomenon may underlie the electrical phenotype of Brugada syndrome, in which hemizygous loss-of-function mutations in Na_V_1.5 produce RV-predominant conduction deficits on ECG. Whereas prior studies attribute this to spatial disparities in repolarizing potassium currents in the heart,^18^ our findings suggest that the RV phenotype of Brugada syndrome results, in part, from reduced Na_V_1.5 and limited compensatory neuronal sodium channel isoforms in the heart, supporting its proposed characterization as a disease of “reduced RVOT conduction reserve.”^19,20^

### Clinical Implications of Non-canonical Sodium Channels in the Heart

Our findings suggest that regional sodium channel isoform expression may modulate both cardiac pathophysiology and drug responses. Heart failure and myocardial ischemia are associated with increased non-inactivating late sodium current, which is thought to contribute to arrhythmia risk and sudden death. Whereas other models attributed increased late currents to changes in Na_V_1.5 expression or function, our data indicate that Na_V_1.8, which displays a rate of fast inactivation far slower than other VGSCs^21^ (except NaV1.9), is enriched in the LV and may mediate pathological late I_Na_. Indeed, Na_V_1.8 has been shown to be enriched in patients with heart failure.^22^

The clear presence of neuronal sodium channel isoforms in atrial myocytes presents possible therapeutic targets for atrial fibrillation, the most common arrhythmia globally,^23^ particularly for patients with limited options for ablation treatments. Similarly, drugs targeting isoforms otherwise limited to the CNS, such as Na_V_1.8, may be effective treatment for arrhythmias associated with structural heart disease, especially in the setting of disease-related changes in sodium channel isoform expression and late sodium currents.^24,25^

Conversely, altered neuronal Na_V_ isoform expression may drive the increased risk associated with sodium channel inhibitors in vulnerable patient populations, as demonstrated by the rise in arrhythmic deaths with flecainide treatment in post-myocardial infarction patients in the CAST trial.^26^ Finally, our findings may inform the safe use novel small molecule inhibitors targeting neuronal sodium channel isoforms (e.g., suzetrigine^14^), as these may potentially carry additional cardiac risk in populations with inherited or acquired forms of heart disease.

### Study Limitations

This study has several strategic weaknesses. We did not assess isoform expression in response to disease states or therapeutic interventions, limiting insight into changes in the isoform landscape during cardiac remodeling. Analyses of isoform composition used inhibitors with incomplete selectivity, resulting in ambiguity in parsing isoforms in certain groups (i.e., Na_V_1.1/1.3 and Na_V_1.2/1.6/1.7). VSTx-3 applied at 1 μM exhibited incomplete blockade of human Na_V_1.8 in cell lines but full inhibition in mouse cardiomyocytes, potentially indicating lower affinity for the human variant. Isoform contributions were estimated across pooled cells and experiments, introducing potential variability. Nonetheless, the sum of the average current contributions closely approximated the total I_Na_, indicating that the measured isoform proportions were complete and mutually exclusive. Finally, findings in mouse cardiomyocytes may not be fully representative of those in humans.

## Methods and Materials

### Sequence of the Na_V_1.5-GX channel

The Na_V_1.5-GX construct was produced by altering the mouse Na_V_1.5 protein sequence from 1556– PEKVNILAKINLLFVAIFTGECIVKMAALRHYYFTNSWNIFDFVVVILSIVGTVL SDIIQKYFF–1620 to 1556– QHMTEVLYWINVVFIILFTGECVLKLISLRHYYFTVGWNIFD FVVVIISIVGMFLADLIETYFV–1620 (**Fig. S1**), which incorporates the GX binding site from human Na_V_1.7. Human and mouse sequences are identical in this region. The region of interest is represented on the structure of rat Na_V_1.5 (**Fig. 1A**, PDB ID: 7FBS).^27^

### Animals

Mice were housed in a temperature-controlled environment with a 12-hr light/dark cycle. Food and water were provided *ad libitum*. Chow containing GX-201 was prepared in DietGel 76A (w/ Animal Protein) (#72-07-5022, ClearH2O, Westbrook, ME). The DietGel was liquified at 60°C, and 0.1 mL of filter-sterilized DMSO containing 612 μg of GX-201 (2.5 mg/kg/day) was injected through the foil lid. The puncture was sealed with tape and the cups were shaken manually for 1-2 min to mix. The cups were stored at 4°C until use. Fresh chow cups were provided daily as the sole source of food. All studies were reviewed and approved by the University of Iowa Institutional Animal Care and Use Committees.

### Generation of the Na_V_1.5-GX mouse

The utilized methods were previous published.^28^ C57BL/6J mice were purchased from Jackson Labs (000664; Bar Harbor, ME). Male mice older than 8 weeks were used to breed with 3–5-week-old super-ovulated females to produce zygotes for pronuclear injection. Female ICR (Envigo; Hsc:ICR(CD-1)) mice were used as recipients for embryo transfer. All animals were maintained in a climate-controlled environment at 25°C and a 12-hr light/dark cycle. Animal care and procedures conformed to the standards of the Institutional Animal Care and Use Committee of the Office of Animal Resources at the University of Iowa.

Chemically modified CRISPR-Cas9 crRNAs and CRISPR-Cas9 tracrRNA were purchased from IDT (Alt-R® CRISPR-Cas9 crRNA; Alt-R® CRISPR-Cas9 tracrRNA (Cat# 1072532)). The crRNAs and tracrRNA were suspended in T10E0.1 and combined to 1 μg/μl (∼29.5 μM) final concentration in a 1:2 (μg:μg) ratio. The RNAs were heated at 98°C for 2 min and allowed to cool slowly to 20°C in a thermal cycler. The annealed cr:tracrRNAs were aliquoted to single-use tubes and stored at -80°C. Cas9 nuclease was purchased from IDT (Alt-R® S.p. HiFi Cas9 Nuclease). Cr:tracr:Cas9 ribonucleoprotein complexes were made by combining Cas9 protein and cr:tracrRNA in T10E0.1 (final concentrations: 300 ng/μl (∼1.9 μM) Cas9 protein and 200 ng/μl (∼5.9 μM) cr:tracrRNA). The Cas9 protein and annealed RNAs were incubated at 37°C for 10 minutes. The RNP complexes were combined with single-stranded repair template and incubated an additional 5 minutes at 37°C. The concentrations in the injection mix were 60 ng/μl (∼0.4 μM) Cas9 protein and 10 ng/μl (∼0.3 μM) each (four guides were used) cr:tracrRNA and 20 ng/μl single-stranded repair template. Pronuclear-stage embryos were collected using methods described in.^28^ Embryos were collected in KSOM media (Millipore; MR101D) and washed 3 times to remove cumulous cells. Cas9 RNPs and double-stranded repair template were injected into the pronuclei of the collected zygotes and incubated in KSOM with amino acids at 37°C under 5% CO2 until all zygotes were injected. Fifteen to 25 embryos were immediately implanted into the oviducts of pseudo-pregnant ICR females.

Guides (5’ to 3’):

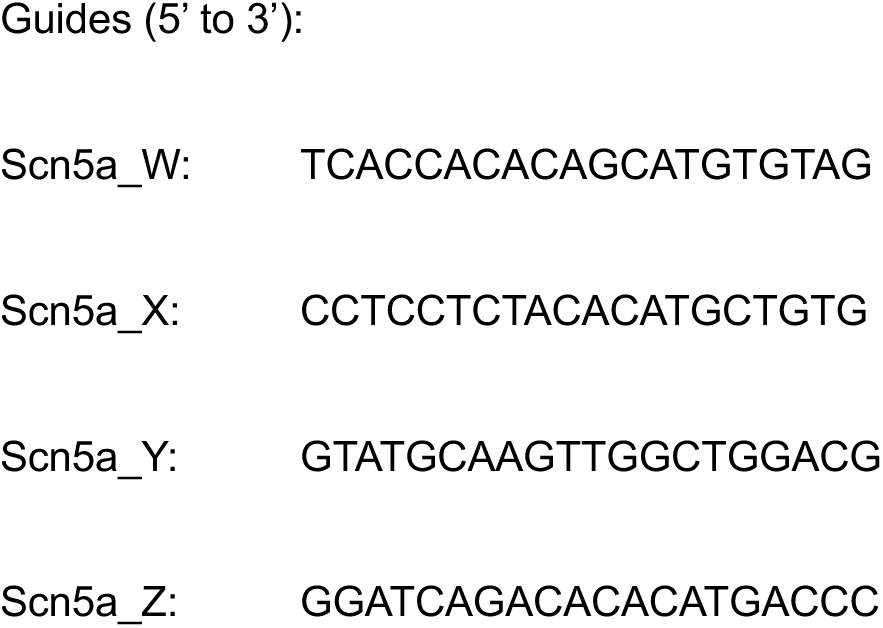

### GX drug selection and synthesis

The GX binding site from Na_V_1.7 was selected for incorporation in Na_V_1.5 because GX compounds are readily available and well-characterized, and Na_V_1.7 has not been significantly implicated in cardiac physiology. We utilized GX-201 (4-[[1-[[2-chloro-5-(trifluoromethyl)phenyl]methyl]-4-piperidinyl]methoxy]-5-cyclopropyl-2-fluoro-N-(methylsulfonyl)benzamide) or MRL5 (5-chloro-2-fluoro-4-(((tetrahydro-1H-pyrrolizin-7a(5H)-yl)methyl)amino)-N-(thiazol-2-yl)benzenesulfonamide) for in vivo studies due to their favorable pharmacokinetics. GX-674 (4-[2-(2-amino-1H-benzimidazol-6-yl)-4-chlorophenoxy]-2,5-difluoro-N-1,2,4-thiadiazol-5-yl-benzenesulfonamide) was used for ex vivo and in vitro studies due to its higher potency.^13^ GX-674, GX-201, and MRL5 were synthesized according to published records^13,29,30^ and authenticated by high-performance liquid chromatography and mass spectroscopy (**Fig. S8**).

### Whole-cell patch clamp electrophysiology

Ionic currents from healthy, quiescent, isolated cardiomyocytes (generated via the Langendorff digestion method) were recorded in whole cell configuration (β=0.1) using Axon Axopatch 200B amplifiers (Molecular Devices). Data were collected and analyzed with pClamp11/Clampfit11 (Molecular Devices) and Origin software (OriginLab). Only well-clamped currents (1 and 20 nA) obtained from cells with access resistance <8 MΩ and >90% series resistance compensation (lag <10 μs) were included.

Recordings were obtained 1-24 hours post-isolation at 20-22°C. The internal solution consisted of 20 mM CsF, 120 mM CsCl, 10 mM NaCl, 10 mM HEPES, 10 mM EGTA, and 5 mM TEA, pH-adjusted to 7.3 with CsOH, 295-305 mOsm adjusted with H_2_O. The main external solution contained 50 mM NaCl, 95 mM CsCl, 1 mM CaCl_2_, 2.5 mM MgCl_2_, and 10 mM HEPES, pH-adjusted to 7.4 with CsOH, 315-325 mOsm adjusted with glucose. The high-Na^+^ external solution used 140 mM NaCl and 5 CsCl and was otherwise identical to the main external solution. Patch pipette tips were 2-3 MΩ with seal resistance >1 GΩ. Data were sampled at 20 kHz and filtered with a lowpass Bessel filter at 10 kHz. Whole-cell capacitance ranged from 75 to 200 pF for ventricular myocytes and 20 to 100 pF for atrial myocytes. Leak currents were subtracted with the P/8 protocol for the current-voltage (IV) protocol but were otherwise not leak subtracted.

In the IV protocol, cells were held at -100 mV and depolarized to -95 to +35 mV for 100 ms in 5 mV increments. In the steady-state inactivation protocol, cells were held at -120 mV, stepped to -110 to +80 mV for 500 ms, hyperpolarized to -100 for 2 ms, and then stepped to a test potential of -20 mV for 20 ms. To assess peak currents during serial inhibitor addition, cells were held at -80 mV, hyperpolarized to -120 mV for 100 ms, and depolarized to -20 mV for 300 ms to elicit peak currents, measured at 1 Hz.

Conductance (*G*) was calculated at each voltage (*V*) using the equation:

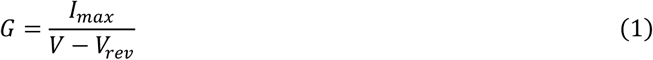

where *I_max_* is the measured peak current amplitude and *V_rev_* is the reversal potential, determined experimentally from the voltage at which net current is zero. Values were normalized to the peak conductance (*G/G_max_*) and fit to a Boltzmann function:

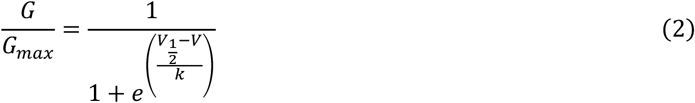

where *V_1/2_* is the voltage of half-maximal activation and *k* is the slope factor.

### Inhibitor application and isoform parsing

Inhibitors were acquired from Alomone Labs (Jerusalem, Israel): VSTx-3 (STT-350), μ-conotoxin GIIIB (C-270) and TTX (T-500). Stock solutions were prepared and stored in -80°C: 1 μM GX-674 in DMSO, 400 μM μ-conotoxin GIIIB in H_2_O, 50 μM TTX in H_2_O, and 1 mM VSTx-3 in H_2_O). For experiments, inhibitors were diluted into 2 ml of external solution to final working concentrations (500 nM GX-674, 200 nM GIIIB, 50 nM TTX, and 1 uM VSTx-3) and mixed gently using a mounted syringe apparatus. Peak currents were monitored until achieving a stable baseline for >1 min before and after addition of each inhibitor. Change in peak current was determined by comparing the average of ≥10 consecutive stable traces before and after each inhibitor addition.

Inhibitors were added sequentially to isolate currents from Na_V_1.5, Na_V_1.2/1.6/1.7, Na_V_1.1/1.3, Na_V_1.4, and Na_V_1.8 as described in the main text. The proportion of current attributed to each Na_V_ isoform was calculated as the fraction of the total baseline current inhibited by each agent. In the ventricles, the proportions of GX-resistant currents (Na_V_1.1/1.3, Na_V_1.4, and Na_V_1.8) were measured in the presence of 500 nM GX-674 and 140 mM Na^+^ external solution and scaled according to the measured fraction of non-Na_V_1.5 and non-Na_V_1.2/1.6/1.7 current. In the atria, the proportions GX-resistant isoforms were measured directly except for Nav1.1/1.3, which was determined by subtracting the average measured fraction of non-Na_V_1.2/1.6/1.7 from the total TTX-sensitive current.

### Inhibitor validation via automated planar patch clamp

Dose-response relationships for GX-674, μ-conotoxin GIIIB, and VSTx-3 were characterized via whole-cell automated planar patch clamp (QPatch Compact, Sophion Bioscience, Ballerup, Denmark) on HEK 293T cell lines (ATCC, Virginia, USA) stably expressing Na_V_1.5-GX or human Na_V_1.3,^31^ Na_V_1.4, Na_V_1.6, or Na_V_1.8 constructs. Na_V_1.8 was modified to include the C-terminal domain of Na_V_1.5 to increase channel expression. Cells were dissociated with Detachin (10-15 minutes at 37°C), triturated in growth media, centrifuged (100 rcf for 4 min), washed with PBS, and resuspended in serum-free media (EX-CELL ACF CHO Medium, MilliPoreSigma, Burlington Massachusetts). External solutions contained: 50 mM NaCl, 4 mM KCl, 95 mM CsCl, 10 mM HEPES, 2 mM CaCl_2_, 1 mM MgCl_2_, 10 mM glucose, pH adjusted to 7.4 with CsOH, 305 mOsm adjusted with sucrose. High-sodium external solution contained 145 mM NaCl and no CsCl but was otherwise identical. Internal solution contained: 140 mM CsF, 10 mM NaCl, 10 mM HEPES, 1 mM EGTA, pH adjusted to 7.3 with CsOH, 320 mOsm adjusted with sucrose.

Cells were recorded in 50 mM external Na^+^ (Nav1.3) or 145 mM external Na^+^ (other isoforms) using the same voltage protocol as in cardiomyocyte experiments. Inhibitors were diluted to working concentrations in external solution and applied sequentially at increasing concentrations. Peak currents were averaged from ≥10 consecutive stable after each inhibitor addition and normalized to the baseline current. The fractional inhibition (*F*) at each concentration (*C*) was fit to a Hill equation:

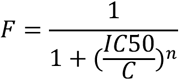

where *IC50* is the concentration of half-maximal inhibition and *n* is the hill coefficient.

### Isolation of adult mouse ventricular cardiomyocytes

Isolated cardiomyocytes were obtained from adult mice between 3 and 6 months of age using the Langendorff perfusion and enzymatic digestion method as previously described,^32^ modified to a constant perfusion pressure configuration. The mice were anesthetized via IP injection with sterile 1.2% Avertin (2,2,2-tribromoethanol and 2-methyl-2-butanol) in PBS at 0.25 ml per 10 g body weight. Heparin was administered simultaneously at 10,000 U/kg. Isolated hearts were mounted on a Langendorff perfusion apparatus (Radnoti Glass Technology, Monrovia, CA, USA, #130107) and perfused with a calcium-free Tyrode’s solution (143 mM NaCl, 5.4 mM KCl, 0.5 mM NaH_2_PO_4_, 22 mM NaHCO_3_, 11 mM glucose, 0.5 mM MgCl_2_, 5 mM butanedione monoxime, pH adjusted to 7.4 with NaOH) at 37°C. Perfusion solutions were bubbled and equilibrated with 5% CO_2_ with 95% O_2_. After 5 minutes, hearts were perfused with 95mg/ml Collagenase Type II (Worthington) and 5mg/ml Collagenase Type IV (Worthington) in calcium-free Tyrode’s until adequately digested (typically 8 to 30 min). The heart was flushed with 3 mL of KB solution (70 mM KOH, 40 mM KCl, 50 mM potassium L-glutamate, 10 mM HEPES, 20 mM taurine, 20 mM KH_2_PO_4_, 10 mM glucose, 3 mM MgCl_2_, 0.5 mM EGTA, pH adjusted to 7.3 with KOH). Hearts were then dissected in KB solution to separate the left ventricle free wall, right ventricle free wall, and atria, which were then dissociated with tweezers and gentle trituration. Isolated cardiomyocytes were allowed to sediment by gravity, resuspended in KB, and stored at room temperature for 1 hour prior to patch clamp experiments.

### Surface electrocardiograms (ECG) and echocardiography

Mouse ECGs and transthoracic echocardiograms were performed as previously described.^33^ ECGs were performed on isoflurane-anesthetized animals with subcutaneous bipolar electrodes in positions corresponding to human ECG leads I, II, III, and modified V. Five to ten consecutive intervals were signal-averaged to improve accuracy. Echocardiography in the long- and short-axis was performed on mice sedated with 10 µL midazolam.

### Immunoblotting

For total protein, mouse heart samples were pulverized under liquid nitrogen and lysed in modified RIPA buffer (50 mM Tris HCI, 150 mM NaCl, 1 mM EDTA, 1% NP-40, 0.1% SDS, 1 mM DTT, 1 mM PMSF, and protease inhibitor cocktail (Promega, Madison, WI). The lysate was clarified by centrifugation at 16,000 rcf for 15 minutes at 4°C.

Protein concentrations were determined via Bradford assay (BioRad Protein Assay) with the Synergy H1 Microplate Reader (Biotek, Winooski, VT). Seventy μg of whole cell lysate were carried out by SDS-PAGE using NuPAGE 4% to 12% Bis-Tris Gel (ThermoFisher, Waltham, MA) ran at 50 mV for 15 minutes, then increased to 80 mV for 2.5 hours in a 1x Running buffer solution (20x MOPS SDS Running Buffer, ThermoFisher). The gel was transferred to PVDF membrane in 1x Transfer buffer solution (10x Tris/Glycine buffer (Biorad, Hercules, CA), 10% methanol, 0.01% SDS) at 35 mV and 4°C overnight.

Membranes were then incubated in 5% nonfat dried milk in Tris-buffered saline/Tween 20 (1x TBS, 0.1% Tween 20) for 30 minutes at room temperature, probed with primary polyclonal antibody for Na_V_1.5. (#ASC-005, Alomone Labs, Jerusalem, Israel) diluted to 1:500 in 5% nonfat dried milk at 4°C overnight, washed 3 times in 1x TBST for 10 minutes, probed with peroxidase-conjugated secondary antibody (Rockland Immunochemicals, Limerick, PA) diluted to 1:5,000 in 5% nonfat dried milk, and washed again in 1x TBST 3 times for 10 minutes. Chemiluminescent signal was detected using a SuperSignal West Dura Extended Duration Substrate (ThermoFisher, Waltham, MA) and imaged with ChemiDoc XRS Gel Imaging System (BioRad, Hercules, CA). Results were normalized to GAPDH.

### Serum Concentration of GX-201

Blood samples (>200 μl) were obtained from mice via cheek bleed or cardiac puncture. The blood was immediately centrifuged at 2,000 rcf for fifteen minutes, and the serum supernatant was transferred and stored at -80°C until assay. Serum samples were lyophilized and placed in 2 ml ceramic bead tubes (Omni International #19-627) with an empty tube as blank control. Extraction was done with 100% Acetonitrile at 18 μl/mg of plasma. Bead tubes were placed on the Bead Ruptor Elite at 6.45 m/s, 01 Cycles, 30 s Time, and 0 s Dwell Time. After homogenization, samples were chilled at -20°C for 1 hour while rotating. Samples were dried and reconstituted in mobile phase buffer (1:1 water:acetonitrile) and analyzed on a Waters XEVO TQ-S Cronos triple quadrupole mass spectrometer with a Waters Acquity H-Class UPLC. The LC column used was a Thermo Hypersil Gold (2.1 x 50 mm, 1.9 μm) column. The mobile phases were 0.1% Formic Acid in Water (A) and Acetonitrile with 0.1% Formic Acid (Fisher Optima LC-MS grade solvents). The flow rate was 0.2 ml/min, and the column was held at 30°C. The gradient started at 35% B and increased linearly over 5.0 minutes to 90% B where it was held for 3 minutes before re-equilibration. An injection of 5 µl was used for both samples and standards. For the LC-MS/MS detection, positive electrospray ionization was used with multiple reaction monitoring (MRM) for detection. The MRM precursor to product ion transitions monitored for GX-201 were 563.1 to 192.7 and 290.1, with collision energies set at 48 and 32 eV, respectively. The MRM transition monitored for MRL5 (internal standard) were 430.97 to 110.02 and 124.07, with collision energies 22 and 24 eV, respectively. Waters MassLynx 4.2 software was used to collect the data, and TargetLynx was used for quantitative analysis.

### ECG Telemetry

ECG radiotelemeters (Data Sciences International, New Brighton, MN, USA: #TA11-ETA-F10) were surgically implanted seven days before the experiment as previously described.^34^ Data was collected at 1 kHz and subsequently filtered with a 4^th^ order Butterworth filter from 30-100Hz for QRS detection. The QRS detection algorithm for this study is based on the XQRS package,^35,36^ a modified implementation of the WFDB algorithm for the Pan & Tompkins method.^37^ XQRS replaces the Pan & Tompkins derivative filter, squaring, and moving window integration with a single-step filter using the Ricker wavelet, followed by squaring the output. For this study, the XQRS algorithm was further adapted to account for mouse physiology by adjusting configuration and filter parameters and omitting the T-wave detection step. Peak detection was based on 70 ms windowing. The final output of the algorithm was a set of indices corresponding to detected QRS complexes which were used for subsequent analysis of R-R intervals and other cardiac metrics. R-R intervals were binned every 1 ms from 60-300 ms. Areas of noisy data were discarded based on the thresholded lookback function, with each telemetry monitoring period containing >16 hours of interpretable data. Telemetry was recorded prior to any drug administration, on days 1, 2, and 3 after drug administration, and again on days 1 and 3 of the washout period with control diets.

Classification of tachy- and brady-arrhythmias was first done automatically by identifying the R-R interval that corresponded to the longest 1.0% of R-R intervals (bradycardia threshold) and the shortest 0.1% of R-R intervals (tachycardia threshold) in the baseline telemetry recording for each individual mouse.

Baseline telemetry tracings were manually reviewed by a Cardiologist to verify normal cardiac conduction. Mice were considered to have bradyarrhythmia if the fraction of R-R intervals that were longer than the bradycardia R-R threshold was >2.5% and considered to have tachycardia if the number of R-R intervals that were shorter than the tachycardia R-R threshold was >1.0%. Tachycardia and bradycardia events were manually reviewed by a board-certified cardiologist to verify arrhythmia and exclude sinus tachycardia or bradycardia.

### RNA Sequencing

Extracted total RNA was sent to CD Genomics for Illumina-based RNA sequencing. Reads were acquired with 150 base-pair, paired-end reads with an average depth of ∼57M reads per sample. Adapter trimmed fastq sequences were aligned using Kallisto^38^ to mouse reference transcriptome GRCm38. Kallisto index and Kallisto quant were run using default settings, with specific input parameters of 100 bootstraps and 16 threads. Differential expression was performed using Sleuth version 0.30.0 aggregating abundance on Ensemble genes, with a two-step Likelihood Ratio Test and Wald Test.^39^ Data are provided in **Table S1**. Raw sequencing reads are deposited into the GEO archive (in progress, release on publication date).

### Optical Mapping

Fluorescence images from the LV and RV free wall of the heart were captured using two CMOS cameras (100×100 pixels, Ultima-L, SciMedia, Japan) with the field of view set to 1.0×1.0 cm^2^ (spatial resolution of 100×100 μm^2^) as previously described.^40^ The sampling rate was set to 2,000 frames per second, and data was analyzed with a custom-built software program developed in Interactive Data Language (Exelis, Inc., Boulder). Hearts were stained with a voltage-sensitive dye, di-4-ANEPPS (Invitrogen, Carlsbad), using 10 µL of stock solution (1 mg/ml of DMSO) delivered through a bubble trap, above the aortic cannula. ECG and perfusion pressure were continuously monitored (PowerLab, ADInstruments, Colorado Springs). The activation and repolarization time points at each site were determined from fluorescence signals by calculating (dF/dt)_max_ and 90% recovery (APD_90_). Data was filtered using a spatial Gaussian filter (7×7 pixel), and first derivatives were calculated using a temporal polynomial filter (3^rd^ order, 13 points). Pixels with low signal-to-noise ratio were determined by (dF/dt)_max_ lower than 3 sample SD from baseline and outliers of pixels determined by Grubbs’ test were removed from analysis. Biphasic upstrokes were detected by the presence of 2 peaks in the first derivative of the action potential data.

### Statistical comparisons

All statistical analyses were performed in using GraphPad Prism v10.4.0 and R v4.5. The statistical tests, sample sizes, and p-values are reported in each figure legend and the main text. For all analyses, a p-value of less than 0.05 was considered statistically significant. Assumptions of normality and equal variance were assessed when necessary. False-discovery rate and multiple-comparisons corrections were applied as described. Error bars represent 1 sample standard deviation unless otherwise stated.

### Data Availability Statement

The data that support the findings of this study are available from the corresponding author upon reasonable request.

## Author Contributions

Conceptualization, C.J.C., D.I., B.L., C.A.A; Methodology, C.J.C., C.A., A.D. B.C.; Investigation C.J.C., C.A., A.D., H.C., J.D., J.G., S.T., H.C., J.Y., B.C., L.G.; Data Analysis and Visualization, C.J.C., C.A., A.D., K.L. H.C., S.T., H.C., J.Y., B.C.; Writing – Original Draft; C.J.C., C.A., A.D.; Writing – Review and Editing; C.J.C., C.A., A.D., B.L., C.A.A.

### Competing Interest Statement

The authors declare no competing interests.

## Acknowledgements

We thank Dr. Alfred George for providing the Na_V_1.3 cell line as well as Dr. Lynn Teesch and the High-Resolution Spectrometry Facility for assistance with mass spectroscopy analysis. Transgenic GX mice were generated at the University of Iowa Genome Editing Core Facility directed by William Paradee, PhD and supported in part by grants from the AHA Transformational Project Award and from the Roy J. and Lucille A. Carver College of Medicine. We thank Dr. William Paradee, Norma Sinclair, Rongbin Guan, and Joanne Schwarting for their technical expertise in generating transgenic GX mice.

## Notes

### Competing Interest Statement

The authors have declared no competing interest.

