## Supplemental file for "Non-canonical Sodium Channel Isoforms Underlie Chamber Specific Cardiac Excitability"

**Supplemental Table 1. mRNA sequencing shows no change in ventricular $Na_V$ expression profile due to the $Na_V1.5$-GX allele.** Average scaled transcripts of $Na_V$ alpha and beta subunits in ventricular tissue from $Na_V1.5^{GX/GX}$ and $Na_V1.5^{WT/WT}$ mice (ND, not detected). Unadjusted p-values from Welch's t-test with adjusted q-value after FDR correction (Benjamini-Hochberg).

| Gene | Average Transcripts | | | | |
|---|---|---|---|---|---|
| | GX/GX | WT/WT | Pooled | p | q |
| Scn1a | ND | ND | | | |
| Scn2a | ND | ND | | | |
| Scn3a | 12.3 | 12.1 | 12.2 | 0.95 | 0.95 |
| Scn4a | 476.0 | 418.2 | 451.2 | 0.46 | 0.91 |
| Scn5a | 7634.6 | 6756.1 | 7258.1 | 0.08 | 0.36 |
| Scn7a | 484.1 | 507.7 | 494.2 | 0.74 | 0.95 |
| Scn8a | ND | ND | | | |
| Scn9a | ND | ND | | | |
| Scn10a | 46.8 | 43.3 | 45.3 | 0.86 | 0.95 |
| Scn11a | ND | ND | | | |
| Scn1b | 387.0 | 330.3 | 362.7 | 0.31 | 0.75 |
| Scn2b | 14.1 | 19.2 | 16.3 | 0.20 | 0.59 |
| Scn3b | ND | ND | | | |
| Scn4b | 1337.0 | 1207.4 | 1281.5 | 0.66 | 0.95 |

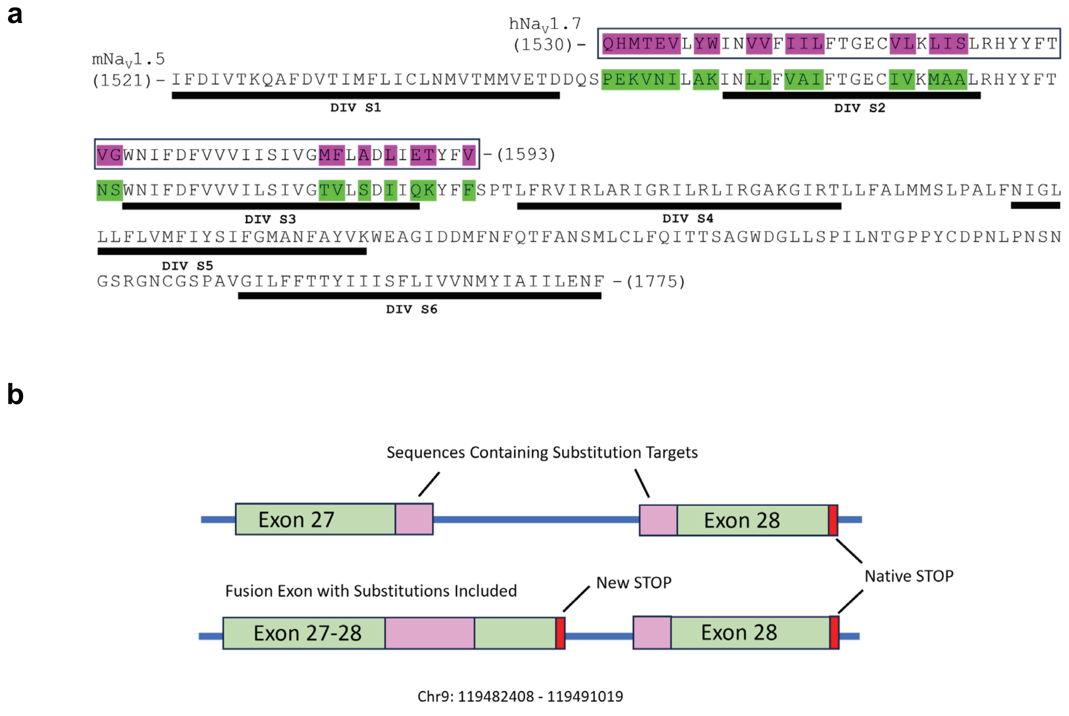


**Supplemental Figure 1: Genetic editing to generate the $Na_V1.5$-GX mice.** (A) Pairwise protein sequence alignment of mouse $Na_V1.5$ and human $Na_V1.7$ showing the region of the domain IV voltage sensor (S2-S3, black box) exchanged to generate the $Na_V1.5$-GX chimeric channel. Non-homologous residues in human $Na_V1.7$ (magenta) or the mouse $Na_V1.5$ background (green) involved in this exchange are highlighted. (B) Editing exons 27 and 28 within the SCN5A genomic locus (Chr9: 119482408-119491019) to encode the $Na_V1.5$-GX construct. A fusion exon containing the chimeric substitutions (pink) and a new stop codon was generated via CRISPR-Cas9 editing. The native exon 28 was left intact.

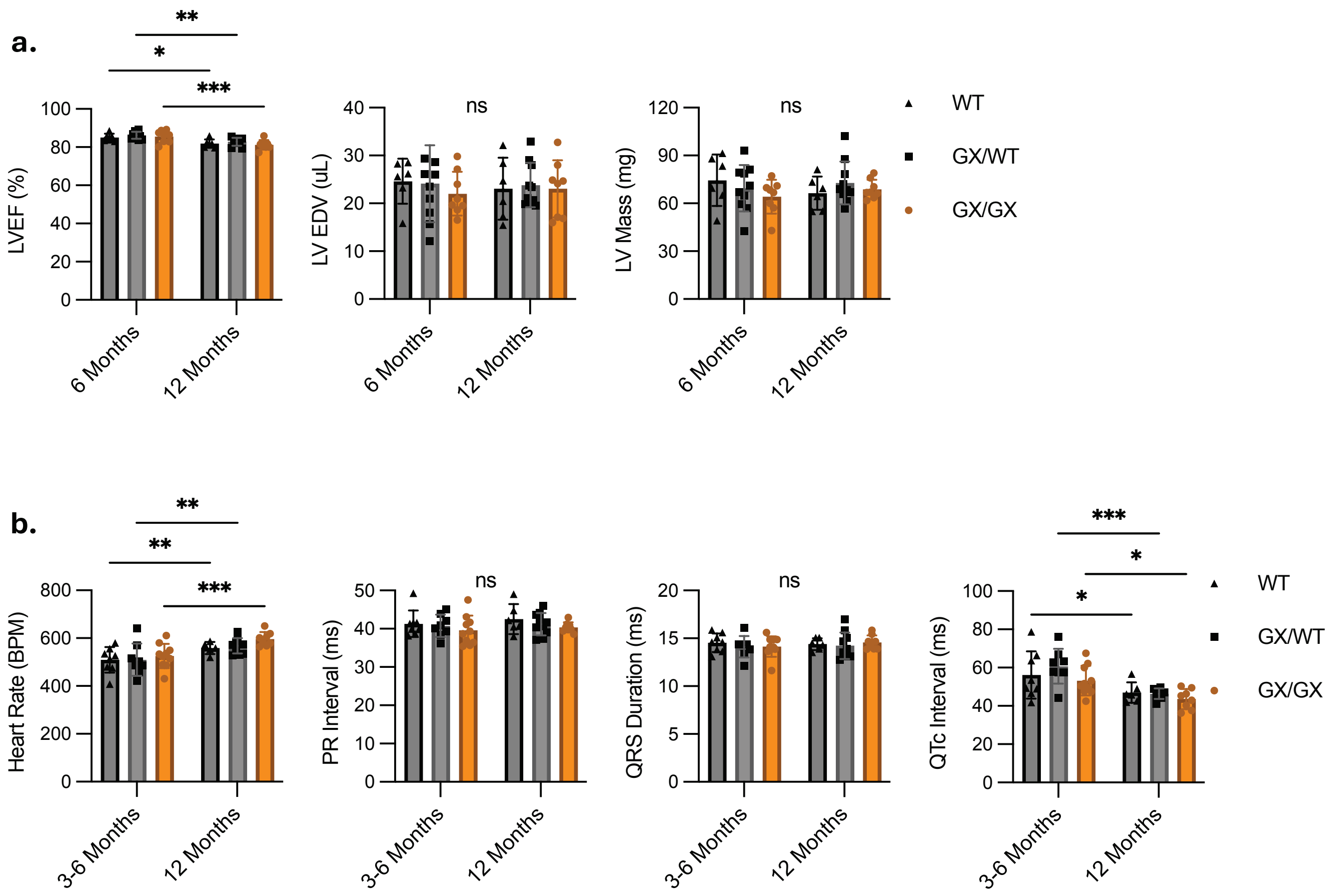


**Supplemental Figure 2: In Vivo Cardiac echocardiography and electrocardiography measurements.** WT (dark gray), heterozygous (light gray) and homozygous $Na_V$1.5-GX (orange) mice were subjected to echocardiography at 6 and 12 months of age (A; LV ejection fraction, LV end diastolic volume, LV mass) and surface ECG measurements between 3 and 6 months of age, and again at 12 months of age (B; heart rate, PR interval, QRS duration, QTc interval). There is a small age dependent decrease in LVEF, otherwise WT, GX/WT, and GX/GX mice did not show any changes in echocardiogram measurements. ECG showed increases in heart rate and decreases in QTc interval between 3-6 months and 12 months but not between groups at each timepoint. No differences were seen in PR interval or QRS duration. Mean±SD. Two-way ANOVA, *$p<0.05$, **$p<0.01$, ***$p<0.001$. n>6 for each echocardiogram measurement, n>5 for each ECG measurement. For echocardiography measurements, cohorts contained equal numbers of males and females.

**a**

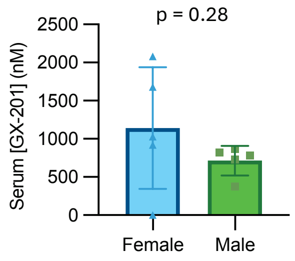


**Supplemental Figure 3:** Serum drug concentrations of mice on oral GX-201. (A) Serum concentrations of GX-201 measured using mass spectroscopy from female (n = 5) and male (n = 5) $Na_V1.5$-GX mice after 3 days of treatment with a 2.5 mg/kg/day oral GX-201 diet. Mean ± SD, two-tailed Student's t-test.

a

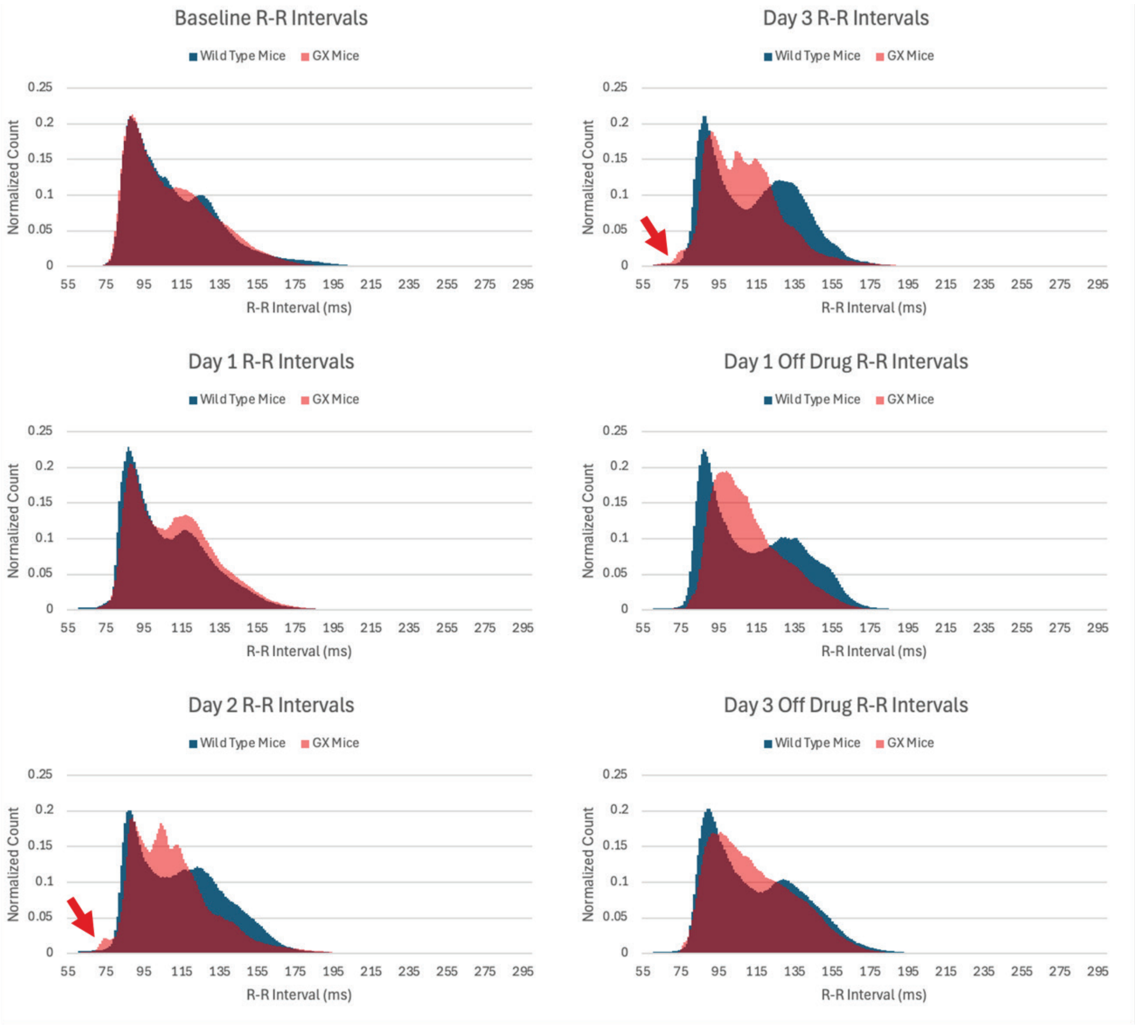


**Supplemental Figure 4: Histograms of R-R Intervals from 24-hour telemetry.** 8 $Na_V1.5$-GX (transparent red) and 7 WT (blue) mice were fed GX-201 chow at a dose of 2.5 mg/kg/day for 3 days with a 3-day washout on a control diet. Telemetry was recorded on Days 0-3, and Days 1 and 3 of washout (Day 0 corresponds to Baseline). Histograms shown are the average for each cohort (n=8 $Na_V1.5$-GX, n=7 WT). R-R intervals were placed in 1 ms bins from 60-300 ms. Histogram counts were normalized using the L2 norm to facilitate overlapped comparison. GX-201 induced conduction changes that resulted in increased bradycardic (long) and tachycardic (short) R-R intervals (red arrows showing tachycardic R-R intervals). These were manually examined for arrhythmia. By day 3 of washout, the histogram approaches the baseline distribution.

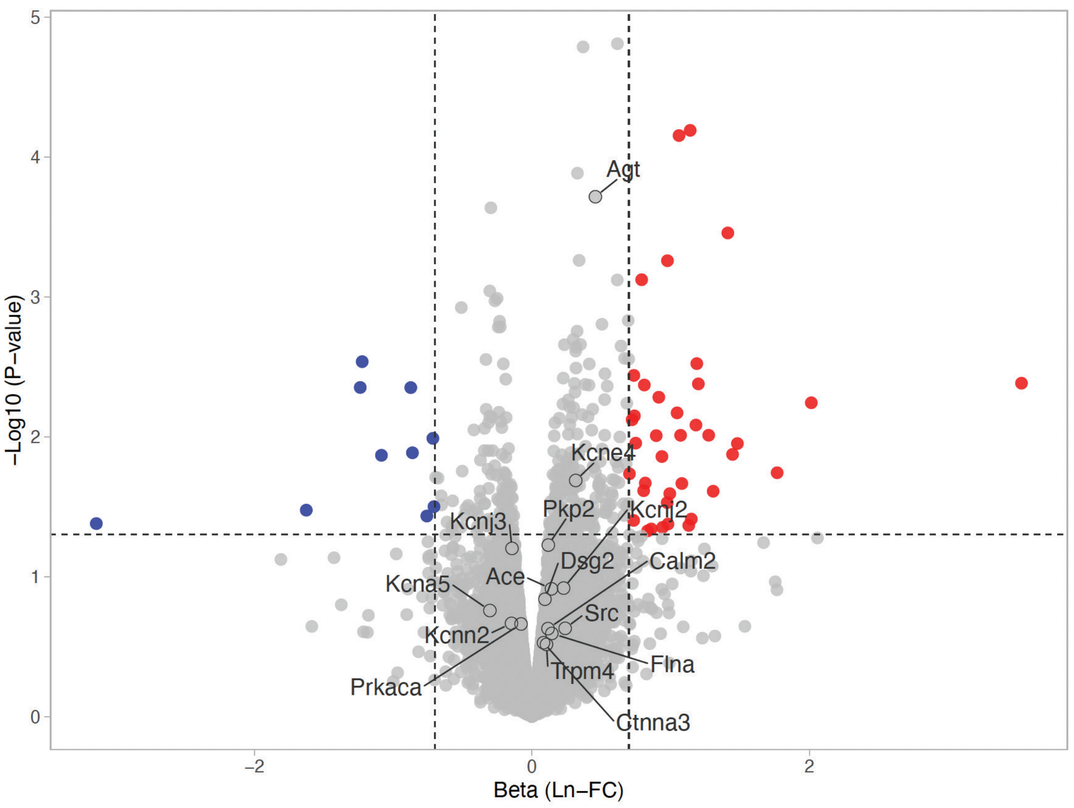


**Supplemental Figure 5: Volcano plot of mRNA-seq data comparing MRL5- and vehicle-treated $Na_V$1.5-GX mice.** Up-regulated (red) and down-regulated (blue) genes based on non-stringent cutoffs (unadjusted P<0.05, beta >0.1) are highlighted. Conduction-related genes are labeled (top 20 ranked by unadjusted P-value) to illustrate that the vast majority are unchanged.

a

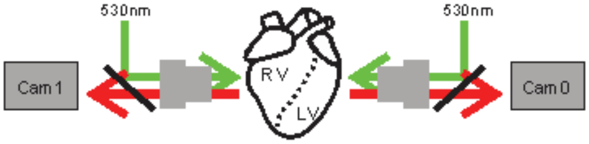


b

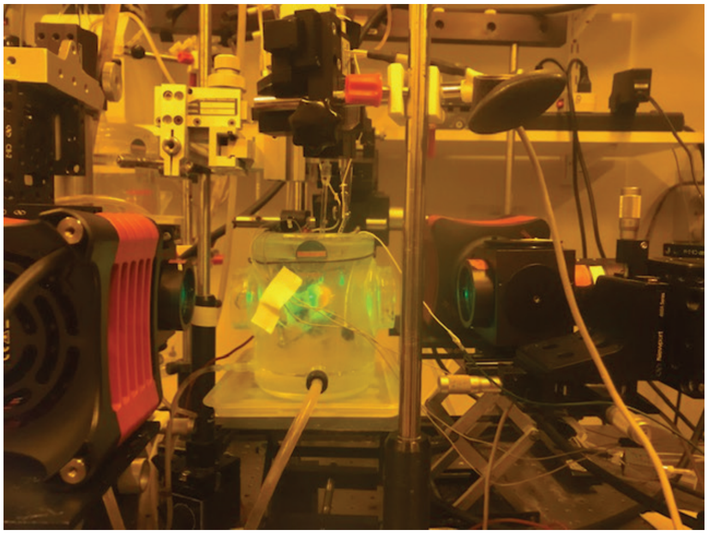

**Supplemental Figure 6: Optical mapping setup.** Schematic (A) and image (B) of the optical mapping system. During imaging, the heart is mounted on a Langendorff perfusion apparatus and perfused at constant pressure with solution containing the voltage-sensitivie dye di-4-ANEPPS. The heart imaging chamber has two windows enabling simultaneous imaging from two cameras. For GX mice, LV and RV freewalls were imaged during stimulation of the central LV freewall with an external electrode. When the atrial tissue (e.g., RA) was stimulated, the heart was positioned such that Cam0 sees the anterior and Cam1 sees the posterior regions of the heart.

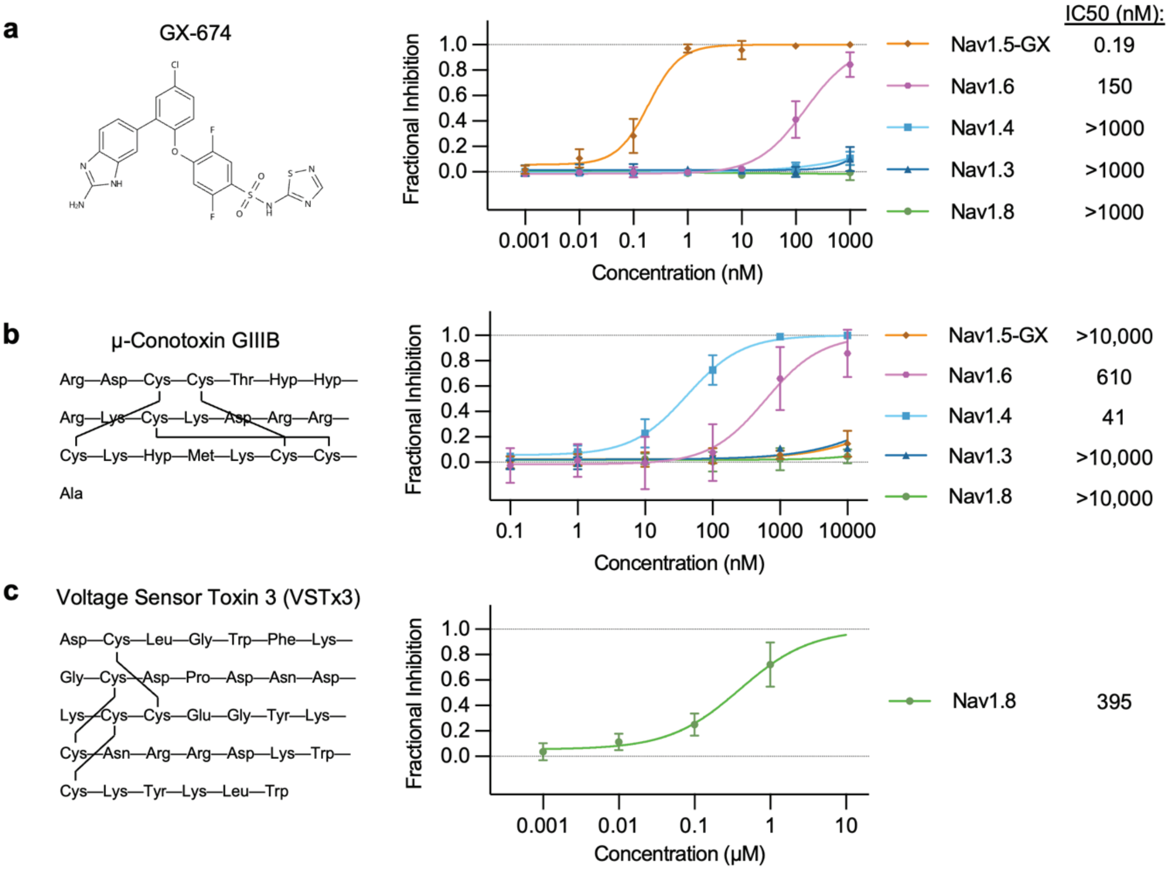


**Supplemental Figure 7: Validation of Nav isoform-selective inhibitors.** (A-C) Structures (left) and dose response curves with Hill equation fit (right) of GX-674 (A), GIIIB (B), and VSTx3 (C) on whole-cell sodium currents of human embryonic kidney cell lines stably expressing $Na_V$1.5-GX (orange diamonds), $Na_V$1.6 (magenta hexagons), $Na_V$1.4 (light blue squares), $Na_V$1.3 (dark blue triangles), or $Na_V$1.8 (green circles) constructs measured via automated planar patch clamp. Cells were held at a potential of -80 mV, hyperpolarized to -120 mV for 100 ms, and depolarized to -20 mV for 300 ms to elicit peak currents at 1 Hz in 50 mM external $Na^+$ ($Na_V$1.3) or 145 mM external $Na^+$ (other isoforms). IC50 values are indicated. Mean ± SD. Data from n ≥ 4 cells for each isoform and inhibitor concentration.

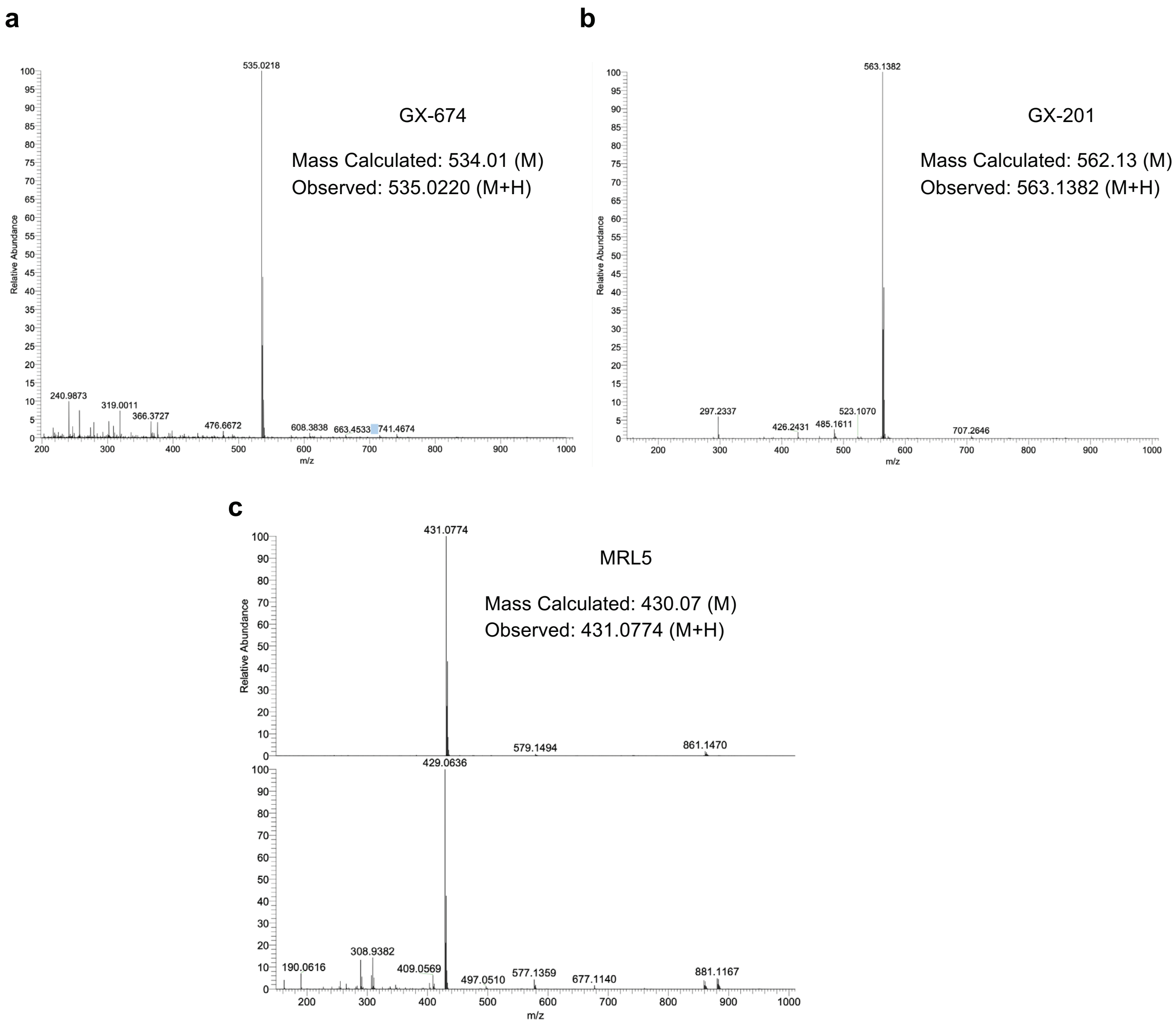


**Supplemental Figure 8: Mass spectroscopy of synthesized GX drugs.** Mass spectra of GX-674 (A), GX-201 (B), and MRL5 (C) show single peaks at the expected masses for each product. Calculated and observed masses of the primary product are indicated.

### Supplemental Video Captions:

**Supplemental Video 1:** Optical mapping of GX/GX hearts at baseline without drugs. Ex vivo hearts stained with voltage-sensitive fluorescent dye (di-4-ANEPPS) from $Na_V1.5^{GX/GX}$ mice were imaged with external stimulation of the left ventricle. The upper panels show the stimulation logic and a schematic of the imaging system. Normal conduction and action potential propagation can be seen through the LV (right) and RV (left) with retrograde conduction through the atria (both) in response to stimulation.

**Supplemental Video 2: Optical mapping of GX/GX hearts in response to GX-674.** Ex vivo hearts stained with voltage-sensitive fluorescent dye (di-4-ANEPPS) from $Na_V1.5^{GX/GX}$ mice were perfused with varying concentrations of GX-674 and imaged with external stimulation of the left ventricle. Ventricular action potential propagation was slowed at 10 nM (left), and nearly absent in the presence of 100 nM GX-674 (right).

**Supplemental Video 3: Optical mapping of chamber specific responses to GX-674 in GX/GX hearts.** Ex vivo hearts stained with voltage-sensitive fluorescent dye (di-4-ANEPPS) from $Na_V1.5^{GX/GX}$ mice were perfused with varying concentrations of GX-674 and imaged with external stimulation of the right atrium. At 30 nM GX-674 there is differential conduction through the RV and LV (left). At 100 nM there is no ventricular excitability and differential conduction between the left and right atria (right).

**Supplemental Video 4: Optical mapping of spontaneous arrhythmia in GX/GX hearts.** Ex vivo hearts stained with voltage-sensitive fluorescent dye (di-4-ANEPPS) from $Na_V1.5^{GX/GX}$ mice were perfused with varying concentrations of GX-674 during spontaneous ventricular activity. The top panels show actional potentials corresponding to spontaneous ventricular activity. 30 nM GX-674 resulted in multiple areas of ectopic ventricular activity propagating ventricular tachycardia (bottom left video). At 10 nM GX-674 there was evidence for reentrant conduction loops in the ventricle also promoting ventricular tachycardia (bottom right video).
